# Developmental dysregulation of chandelier cell excitability in a mouse model of Dravet Syndrome

**DOI:** 10.64898/2026.02.02.702918

**Authors:** Sophie F. Hill, Emiola A. Enakhimion, Elisabetta Furlanis, Min Dai, Brenda Leyva Garcia, Sara Wills, Thien Tran, Gordon Fishell, Yating Wang, Ethan M. Goldberg

**Affiliations:** Division of Neurology, Department of Pediatrics, The Children’s Hospital of Philadelphia, Philadelphia, PA; College of Arts and Sciences, The University of Pennsylvania, Philadelphia, PA; Blavatnik Institute, Department of Neurobiology, Harvard Medical School, Cambridge, MA; Stanley Center for Psychiatric Research, Broad Institute of MIT and Harvard, Boston, MA; Epilepsy NeuroGenetics Initiative, The Children’s Hospital of Philadelphia, Philadelphia, PA; Department of Neuroscience, The University of Pennsylvania, Philadelphia, PA; Department of Neurology, The University of Pennsylvania, Philadelphia, PA

**Keywords:** *Scn1a*, Dravet Syndrome, Chandelier cell, Epilepsy, Sodium channel

## Abstract

Chandelier cells (ChCs) are a rare and highly specialized sub-type of inhibitory neuron in the cerebral cortex that specifically and exclusively form synapses onto the axon initial segment of excitatory neurons and thereby exert powerful control of excitability. We applied a newly developed, cell type-specific enhancer element to label ChCs and assess their electrophysiological and morphological properties across development in Dravet Syndrome (*Scn1a*^*+/-*^ mice), a prominent neurodevelopmental disorder defined by epilepsy and features of autism spectrum disorder. Dravet Syndrome is known to predominantly impact GABAergic interneurons. We found that ChCs from juvenile *Scn1a*^*+/-*^ mice (postnatal day, P18-21) exhibit impaired excitability, and that deficits largely persist in young adulthood (P35-56). These findings are distinct from prior observations in the more common subtype of parvalbumin-positive interneurons, basket cells. We found no differences in the axonal or dendritic morphology of ChCs at either time point. Our results suggest a role for ChCs in the pathogenesis of Dravet Syndrome and constitute the first targeted study of ChC function in a specific genetically-defined neurodevelopmental disorder.

## Introduction

Parvalbumin-expressing, GABAergic inhibitory neurons (PVINs) constitute 40% of all inhibitory neurons in the neocortex (1). Based on morphology, these cells can be sub-divided into two groups: basket (BCs) and chandelier cells (ChCs). BCs make up the bulk of neocortical PVINs and form synapses onto the soma and proximal dendrites of target cells, including pyramidal cells as well as various types of interneurons (1). In contrast, ChCs exhibit exquisite target specificity, exclusively forming synapses onto a discrete subcellular compartment of one target cell type, the axon initial segment (AIS) of pyramidal cells, and thereby control action potential generation (2). These synapses occur in “cartridges” of 4-6 synapses along the length of the AIS (2–4) and confer upon the ChC axon its characteristic “chandelier”-like appearance. A single ChC can contact dozens to hundreds of nearby pyramidal cells (5), and the precise synaptic targeting of AISs endows powerful and direct control over pyramidal cell output (6).

BCs and ChCs can also be distinguished based on their electrophysiological properties. Mature BCs exhibit characteristic fast-spiking behavior, with sustained non-adapting trains of brief action potentials at high firing frequencies exceeding 300 Hz (4). While also considered fast-spiking cells, ChCs exhibit lower maximal firing frequencies (150-200 Hz) (1, 4, 7). Developmentally, ChCs and BCs are derived from the same population of *Nkx2*.*1*^*+*^ progenitor cells in the medial ganglionic eminence (1, 3). However, ChCs are born several gestational days later than BCs in mouse, and ChC electrophysiological properties have been reported to mature more slowly (3, 4).

The primary sodium channel α-subunit isoform expressed in BCs is Nav1.1, encoded by *SCN1A* (8, 9). Heterozygous loss of *SCN1A* causes the neurodevelopmental disorder known as Dravet Syndrome, which is characterized by early-onset, drug-resistant epilepsy; developmental delay and intellectual disability with a formal diagnosis of autism spectrum disorder (ASD) in up to 70% of cases (10); and ataxia (11). In mice, PVIN-specific deletion of *Scn1a* recapitulates many of the key features of Dravet Syndrome (12, 13). We have previously shown that *Scn1a*^*+/-*^ BCs exhibit reduced firing frequency in adolescence (postnatal day (P) 18-21), a deficit that largely recovers by early adulthood (P35-56) (14). However, deficits in synaptic transmission persist (15), which may explain the durable intellectual disability and features of ASD observed in affected individuals.

Until recently, ChCs have been challenging to study due to the difficulty of specifically targeting fluorescent reporters or calcium indicators to these cells. The recent discovery of an enhancer element that can achieve ChC-specific expression has facilitated a more efficient study of ChC function (16).

Here, we applied this enhancer to drive expression of a fluorescent reporter which enabled us to study the potential role of ChCs in the pathogenesis in Dravet Syndrome. We find that *Scn1a*^*+/-*^ ChCs exhibit impaired excitability both at P18-21 and at P35-56. We observed no change in the morphology of *Scn1a*^*+/-*^ ChCs at either time point. These findings demonstrate that all PVINs are impaired in Dravet Syndrome and hence may play a role in disease pathogenesis. The developmental trajectory of this impairment is distinct from that observed in basket cells, suggesting a potential role for ChC dysfunction in the chronic phase of the disorder.

## Methods

### Mice

All experiments were approved by the Institutional Animal Care and Use Committee at the Children’s Hospital of Philadelphia (CHOP) and conducted in accordance with the ethical guidelines of the National Institutes of Health.

Male and female mice were used for all experiments in roughly equal proportions. All mice were on a mixed 50:50 C57/Bl6J:129/SvEvTac genetic background. Experimental animals were generated by crossing 129.textitScn1a^+/-^ mice (Jax #037107) (17) to C57/Bl6J mice (Jax #000664). Litters were weaned between P21-28, and males and females were subsequently housed separately with up to 5 animals per cage. Animals were maintained on a 12/12h light/dark cycle with ad libitum access to food and water.

### Viral injections

On P1, mice were cryoanesthetized on ice for 2 minutes. Virus (AAV.PhP.eB.BiPVe4.dTomato16, 2E12 gc/mL, 1 µL per hemisphere) was injected bilaterally into somatosensory cortex (coordinates relative to bregma: approximately -1.0 mm antero-posterior, ± 2.5 mm medio-lateral, -0.25 mm dorso-ventral) using a 33G Hamilton syringe. Mice were allowed to recover on a heating pad for 10 minutes before being returned to the home cage.

### Acute brain slice preparation

Brain slices were prepared from mice at P18-21 or P35-56. Mice were anesthetized with isoflurane. Older mice (P35-56) were transcardially perfused with ice-cold sucrose cutting solution (in mM: 75 sucrose, 10 glucose, 26 sodium bicarbonate, 2.5 potassium chloride, 1.25 sodium phosphate monobasic, 87 sodium chloride). Brains were removed to ice-cold cutting solution bubbled with 95% O2/5% CO2. Slices were cut on a vibratome (Leica VT1200S) at a thickness of 300 µm. Slices were incubated for 30 mins in cutting solution at 32°C before recording.

### Whole-cell patch clamp recordings

Recordings were performed on a SliceScope electrophysiology rig (Scientifica). Signals were sampled at 100 kHz with a MultiClamp 700B amplifier (Molecular Devices) and digitized with a Digidata 1550B (Molecular Devices). Signals were acquired using pClamp 11 software (Molecular Devices) and low-pass filtered at 10 kHz. All recordings were performed in artificial cerebrospinal fluid (in mM, 10 glucose, 125 sodium chloride, 2.5 potassium chloride, 26 sodium bicarbonate, 1.25 sodium phosphate monobasic, 2 calcium chloride, 1 magnesium sulfate) bubbled with 95% O2/5% CO2 and heated to 32 ± 2°C. Pipettes were pulled on a PC-100 pipette puller (Narishige) to a final tip resistance of 3-6 MΩ. Pipettes were filled with an internal solution (in mM, 130 potassium gluconate, 6.3 potassium chloride, 1 magnesium chloride, 10 HEPES, 0.5 EGTA, 4 ATP-magnesium, 0.4 GTP-sodium, pH 7.4 and osmolarity 285-305 mOsmol/kg) with 0.5% biocytin (Tocris). Liquid junction potential was calculated to be 14.9 mV at 32°C and was not corrected.

All recorded cells were located in primary somatosensory cortex, which was identified based on the presence of characteristic “barrels” with a 4X air objective. Only cells at the layer 1/2 border, where ChCs are known to be concentrated3, were recorded. Slices were visualized under differential interference microscopy, and presence of the dTomato fluorescent tag was used to identify putative ChCs. Cells were excluded if the resting membrane potential was more depolarized than -50 mV or if series resistance exceeded 20 MΩ or changed by more than 20% throughout recording.

After the whole-cell configuration was achieved, cells were allowed to equilibrate for 2 minutes. Two protocols were then applied in current clamp. The first was a one-minute gap-free recording to establish the resting membrane potential. For subsequent recordings, bias current was applied to maintain the membrane potential at -70 mV. The second protocol was a series of 600 ms current steps beginning at -50 pA and increasing to 600 pA in 5 pA intervals with an inter-sweep interval of 2s. This recording was used for analysis of intrinsic properties, as described below.

### Electrophysiology data analysis

All analysis was performed with custom Matlab (version R2023a) scripts and validated manually in ClampFit software (Molecular Devices). Resting membrane potential was defined as the average membrane voltage during the one-minute current-clamp gap-free recording. Input resistance was calculated from the steady-state difference in voltage before and after application of a -50 pA, 600 ms current pulse. Membrane time constant was calculated by fitting a single exponential to the first 200 ms of the response to -50 pA current injection. Membrane sag was calculated as the ratio of the steady-state response to hyperpolarizing current injection to the instantaneous response.

Action potentials (APs) were defined as voltage spikes with minimum slope of 10 mV/ms and minimum amplitude of 30 mV. AP threshold was defined as the point at which the slope of the AP exceeded 10 mV/ms. Maximum steady-state firing frequency was calculated as the highest number of APs during a current injection divided by the duration of the current injection. Maximum instantaneous firing frequency was calculated as the reciprocal of the shortest inter-spike interval (defined as the time difference between AP threshold from one spike to the next). Spike frequency adaptation was the ratio of the first and last inter-spike interval in the first sweep that exceeded 35 APs.

### Tissue clearing and immunostaining

CUBIC tissue clearing (18) was performed with a modified protocol, as follows. On day 1, slices were frozen on dry ice and thawed, then washed 3×10 minutes at room temperature in 0.1M phosphate buffer (0.1M sodium phosphate dibasic brought to pH 7.4 with 0.1M sodium phosphate monobasic). Slices were incubated in CUBIC reagent 1 (for 50 mL: 14.4 mL water, 12.5 g urea, 15.4 mL 80% N,N,N,N-Tetrakis(2-Hydroxypropyl)ethylenediamine (quadrol) in water, 7.1 mL Triton X-100) overnight at room temperature. On day 2, slices were washed 6×15 minutes in 0.1M phosphate buffer at room temperature, then incubated in blocking buffer (5% normal goat serum, 0.3% Triton X-100 in 0.1M phosphate buffer) overnight at 4°C. On day 3, slices were transferred to primary antibody (guinea pig anti-AnkG, Synaptic Systems #386 005, 1:500) diluted in blocking buffer and incubated overnight at 4°C. On day 4, slices were washed 6×15 minutes in 0.1M phosphate buffer at room temperature, then transferred to secondary antibody (goat anti-guinea pig 633, Invitrogen #A21105, 1:500) and streptavidin-488 (Molecular Probes #S32354, 10 ug/mL) diluted in blocking buffer and incubated overnight at 4°C. On day 5, slices were washed 6×15 minutes in 0.1M phosphate buffer at room temperature, then transferred to CUBIC reagent 2 (for 50 mL, 7.5 mL water, 12.5 g urea, 25 g sucrose, 5 g triethanolamine) and incubated overnight at room temperature. On day 6, slices were mounted in CUBIC reagent 2. Imaging was performed within one week.

### Biocytin filling and fixation

Biocytin-filled cells were visualized and imaged on a Leica SP8 confocal microscope. In each slice, the filled cell was located by eye under epifluorescence. A 300 µm z-stack with 1 µm z-steps spanning the entire slice was then acquired. Morphology was evaluated by trained observers blind to genotype. Only cells confirmed to exhibit biocytin-filled cartridges overlapping with AnkG-labeled AISs were included in the study.

For detailed analysis of morphology, a subset of well-filled confirmed ChCs (3 cells per genotype and time point, 12 total) were reconstructed in Neurolucida360 software (MBF Biosciences). All reconstructions were performed manually by an experienced user blind to age and genotype. Quantification was performed in Neurolucida Explorer software (MBF Biosciences).

## Results

### Specificity of ChC viral enhancer

The specificity of the ChC viral enhancer and other interneuron subtype-specific enhancers depends on the route of administration and the age of injection (16). In general, lower injection titers and older age at administration are associated with higher specificity (16). In order to achieve detectable labeling by P18, our earliest time point for electrophysiology, we administered AAV.PhP.eB.BiPVe4.dTomato at P1 via intraparenchymal injection. Each cell was filled with biocytin to confirm ChC morphology based on overlap of biocytin-filled cartridges with AnkG AIS immunolabeling.

Using this method, we observed that 43% of dTomato+ cells at the border of layers 1 and 2 in somatosensory cortex were truly ChCs (Fig 1A). This proportion was not affected by age or genotype (Fig 1B). The remaining dTomato+ recorded cells exhibited fast spiking (16%), regular spiking (24%), or other (17%) firing patterns in roughly equal proportions (Fig 1A).

**Fig. 1.**
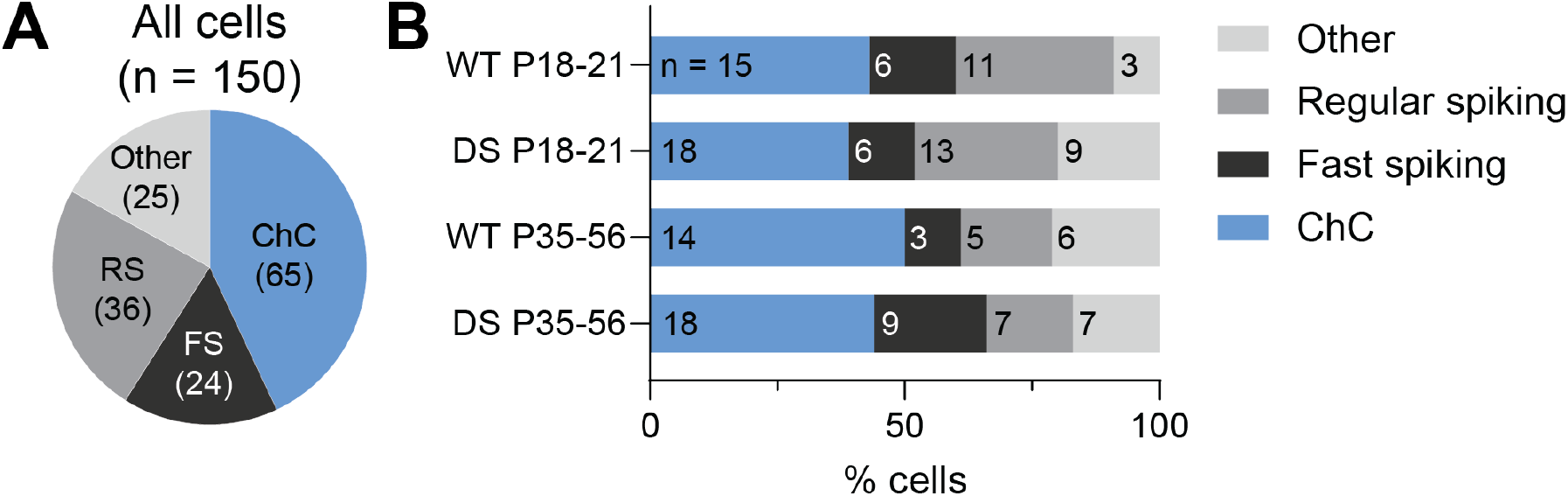
Specificity of chandelier cell enhancer virus. On P1, AAV.PhP.eB.BiPVe4.dTomato was administered to WT and DS mice via intraparenchymal injection. **A**, Proportion of all recorded dTomato+ cells that were confirmed ChCs (40%). Firing pattern of the non-ChCs is shown: FS, fast spiking confirmed non-ChC; RS, regular spiking; or other. **B**, Breakdown by age and genotype. In all conditions, roughly 40% of recorded dTomato^+^ cells were ChCs.

### ChCs are impaired in developing Dravet Syndrome mice

We performed targeted electrophysiology in dTomato^+^ cells at the layer 1/2 border in primary somatosensory cortex (Fig 2A). At P18-21, we observed a clear reduction in the excitability of anatomically confirmed ChCs in *Scn1a*^*+/-*^ mice compared to WT (Fig 2B-D). *Scn1a*^*+/-*^ ChCs exhibited lower maximal steady-state firing frequency (WT: 175 Hz ± 62, DS: 114 ± 20, p = 0.0095; Fig 2, Supplementary Table 1). The first APs at rheobase in *Scn1a*^*+/-*^ ChCs were wider (WT: 0.37 ± 0.10 ms half-width, DS: 0.47 ± 0.06, p = 0.0244) and exhibited reduced upstroke (WT: 351 mV/ms ± 65, DS: 282 ± 55, p = 0.0077) and downstroke velocity (WT: -185 mV/ms ± 67, DS: -135 ± 32, p = 0.0427; Fig 2J-M, Supplementary Table 1). We also observed reduced input resistance in *Scn1a*^*+/-*^ ChCs compared to WT (WT: 142 MΩ ± 49, DS: 107 ± 24, p = 0.0488). These findings are consistent with loss of sodium current as well as with observations in *Scn1a*^*+/-*^ PVIN basket cells at P18-21 (14), suggesting that Nav1.1 plays a similar role in both basket and chandelier cell PVINs at this developmental time point.

**Fig. 2.**
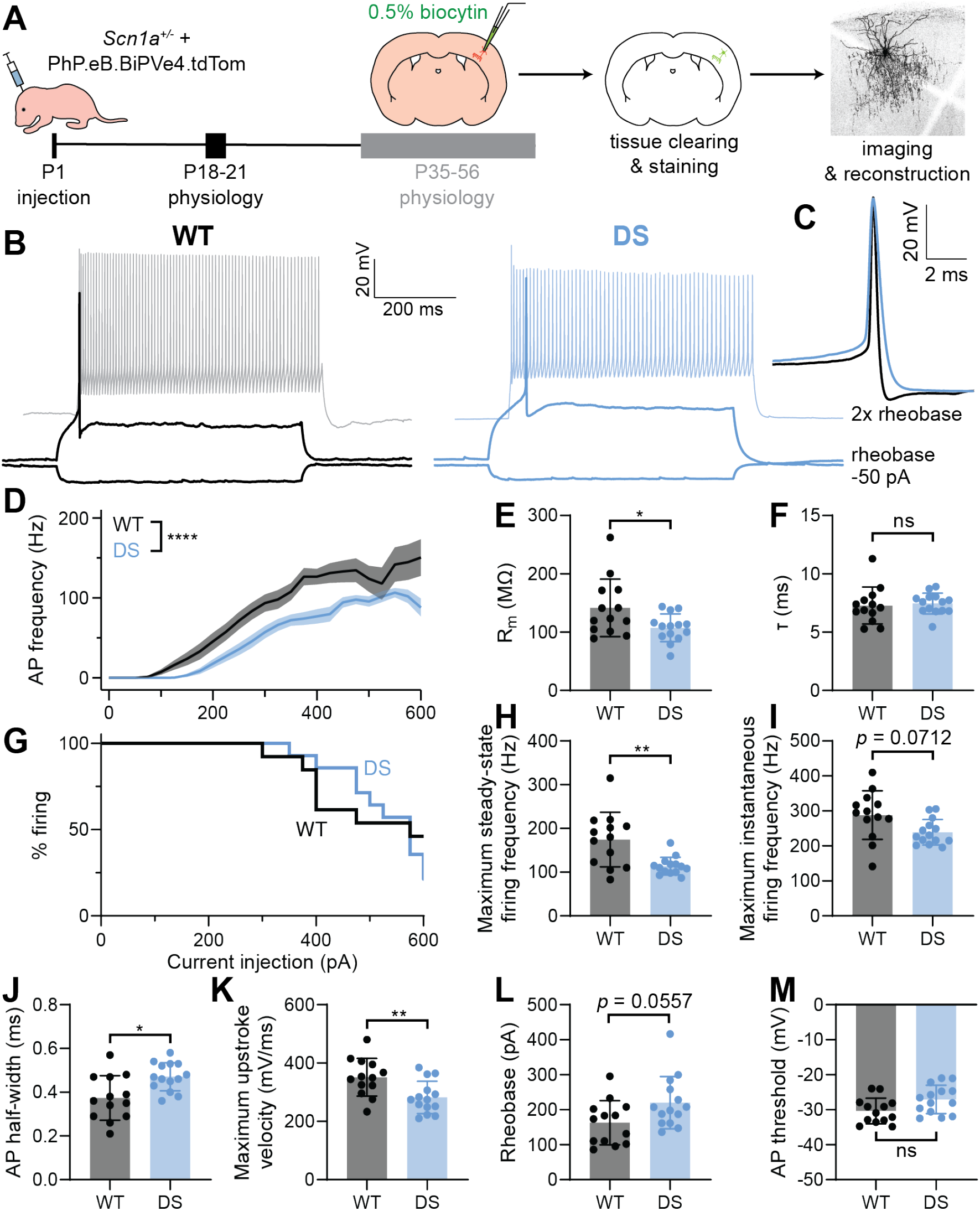
Dravet Syndrome chandelier cells exhibit reduced firing frequency in adolescence. **A**, Mice were injected on P1 with AAV-PhP.eB.BiPVe4.dTomato to fluorescently label chandelier cells (ChCs). At P18-21, acute brain slices were prepared and dTomato^+^ cells in somatosensory cortex L1/2 were recorded and filled with biocytin. Slices were fixed and cleared, and cells were imaged on a confocal microscope to confirm morphology. **B**, Example current-clamp traces of P18-21 WT and DS chandelier cells. Responses to -50 pA, rheobase, and 2x rheobase stimulation are shown. **C**, Larger view of the first action potential (AP) at rheobase from the cells in B. **D**, Input-output curves of WT and DS ChCs (generalized linear mixed-effects model, genotype p < 0.0001, current injection p < 0.0001, interaction p < 0.0001). **E**, Input resistance. **F**, Membrane time constant. **G**, Survival analysis of WT and DS ChCs. Cells were excluded when the dropped below their maximum firing frequency (Mantel-Cox log-rank test, p = 0.5262). **H**, Maximum steady-state firing frequency. **I**, Maximum instantaneous firing frequency. **J-M**, Properties of the first AP at rheobase. Asterisks and p-values indicate the significance of generalized linear mixed-effects models that account for the effects of age, sex, and recording multiple cells from the same animal. * indicates p < 0.05; ** indicates p < 0.01. ns, not significant.

### ChC excitability remains impaired in young adulthood in Dravet Syndrome

In contrast to neocortical basket cells, the excitability of *Scn1a*^*+/-*^ ChCs largely remained impaired at P35-56 (Fig 3). AP frequency was reduced at current injections between 200-400 pA (Fig 3D). This effect was largely driven by a subset of *Scn1a*^*+/-*^ ChCs that exhibited premature depolarization block at intermediate current injections (Fig 3G).

**Fig. 3.**
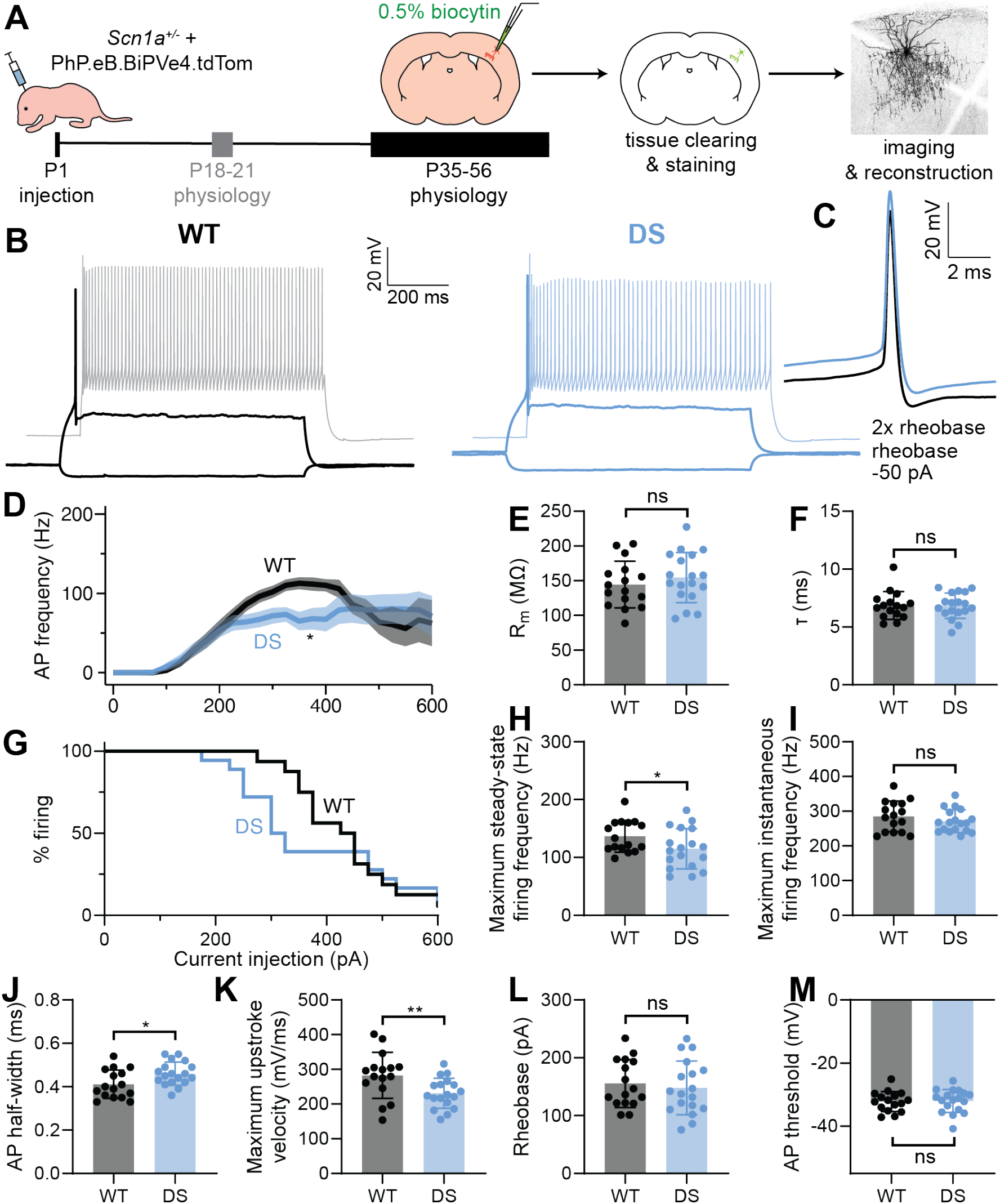
Dravet Syndrome chandelier cells remain impaired in early adulthood. **A**, Mice were injected on P1 with AAV-PhP.eB.BiPVe4.dTomato to fluorescently label chandelier cells (ChCs). At P35-56, acute brain slices were prepared and dTomato^+^ cells in somatosensory cortex L1/2 were recorded and filled with biocytin. Slices were fixed and cleared, and cells were imaged on a confocal microscope to confirm morphology. **B**, Example current-clamp traces of P35-56 WT and DS chandelier cells. Responses to -50 pA, rheobase, and 2x rheobase stimulation are shown. **C**, Larger view of the first action potential (AP) at rheobase from the cells in B. **D**, Input-output curves of WT and DS ChCs (generalized linear mixed-effects model, genotype p = 0.1775, current injection p < 0.0001, interaction p = 0.0036; asterisks show significance of post-hoc Sidak’s multiple comparison tests). **E**, Input resistance. **F**, Membrane time constant. **G**, Survival analysis of WT and DS ChCs. Cells were excluded when the dropped below their maximum firing frequency (Mantel-Cox log-rank test, p = 0.3310). **H**, Maximum steady-state firing frequency. **I**, Maximum instantaneous firing frequency. **J-M**, Properties of the first AP at rheobase. Asterisks and p-values indicate the significance of generalized linear mixed-effects models that account for the effects of age, sex, and recording multiple cells from the same animal. * indicates p < 0.05; ** indicates p < 0.01. ns, not significant.

Similarly, many of the changes in ChC intrinsic properties identified at P18-21 remain abnormal at P35-56, although the magnitude of the changes is slightly reduced (Fig 3, Supplementary Table 2). The maximum steady-state firing frequency is reduced in *Scn1a*^*+/-*^ compared to WT ChCs at P35-56 (WT: 137 Hz ± 27, DS: 115 ± 35, p = 0.0482), although there is no difference in input resistance (WT: 144 MΩ ± 34, DS: 155 ± 36, p = 0.3952). The upstroke (WT: 283 mV/ms ± 66, DS: 231 ± 43, p = 0.0088) and downstroke velocity (WT: -164 mV/ms ± 47, DS: -135 ± 32, p = 0.0360) remain reduced in *Scn1a*^*+/-*^ ChCs, and the AP half-width (WT: 0.41 ms ± 0.07, DS: 0.46 ± 0.05, p = 0.0195) remains lengthened (Fig 3, Supplementary Table 2).

Because ChCs mature later than BCs, we hypothesized that the compensatory process that restores BC excitability at P35-56 may not yet be complete in ChCs. Thus, ChCs recorded around P35 would be predicted to be more impaired than ChCs recorded around P56. We tested this possibility by examining the relationship between age and electrophysiological properties. We found no significant correlation of age with maximum steady-state firing frequency, AP half-width, or maximum upstroke velocity in either genotype within the P35-56 age range (Fig 4). Thus, it is unlikely that the lack of electrophysiological normalization in P35-56 *Scn1a*^*+/-*^ ChCs is due to the delayed maturation of ChCs relative to BCs.

**Fig. 4.**
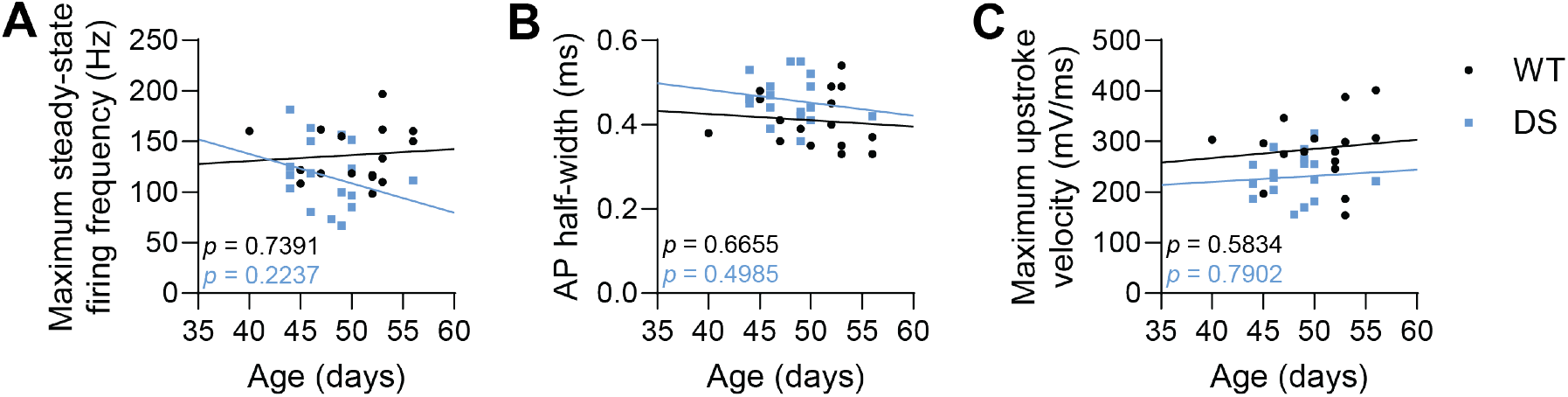
Age does not correlate with chandelier cell impairment at P35-56. Correlation of maximum steady-state firing frequency (**A**), action potential (AP) half-width (**B**), and maximum upstroke velocity (**C**) with age in the P35-56 range. P values indicate significance of slopes from regression lines derived from generalized linear mixed effects models that includes a term for interaction between genotype and age; color indicates genotype.

### ChC morphology is not altered in Dravet Syndrome

To assess possible changes in ChC morphology, we performed full neuronal reconstructions for three ChCs of each age and genotype (Fig 5A). We quantified the total number of axonal branches (a proxy for the number of cartridges), maximal axonal branch order, total axonal length, and length of the axon initial segment (Fig 5B-E). We identified no change in any of these parameters between WT and *Scn1a*^*+/-*^ ChCs except that the total axonal length was slightly higher in *Scn1a*^*+/-*^ ChCs at P18-21 (WT: 10.6 mm ± 2.4, DS: 13.4 ± 1.4, p = 0.0224; Fig 5D). We also performed Sholl analysis to compare the spatial distribution of axonal branching between WT and *Scn1a*^*+/-*^ ChCs. We identified no changes in the number of intersections at P18-21 or P35-56 (Fig 5F-G). We quantified the total number of dendritic branches, maximal dendritic branch order, and total dendrite length (Fig 5H-J). There was no difference in any dendritic metric between WT and *Scn1a*^*+/-*^ ChCs. Sholl analysis of dendrites also revealed no change in the spatial distribution of dendritic branches at P18-21 or P35-56 (Fig 5K-L). From these data, we conclude that the morphology of ChCs is not altered by haploinsufficiency of *Scn1a*.

**Fig. 5.**
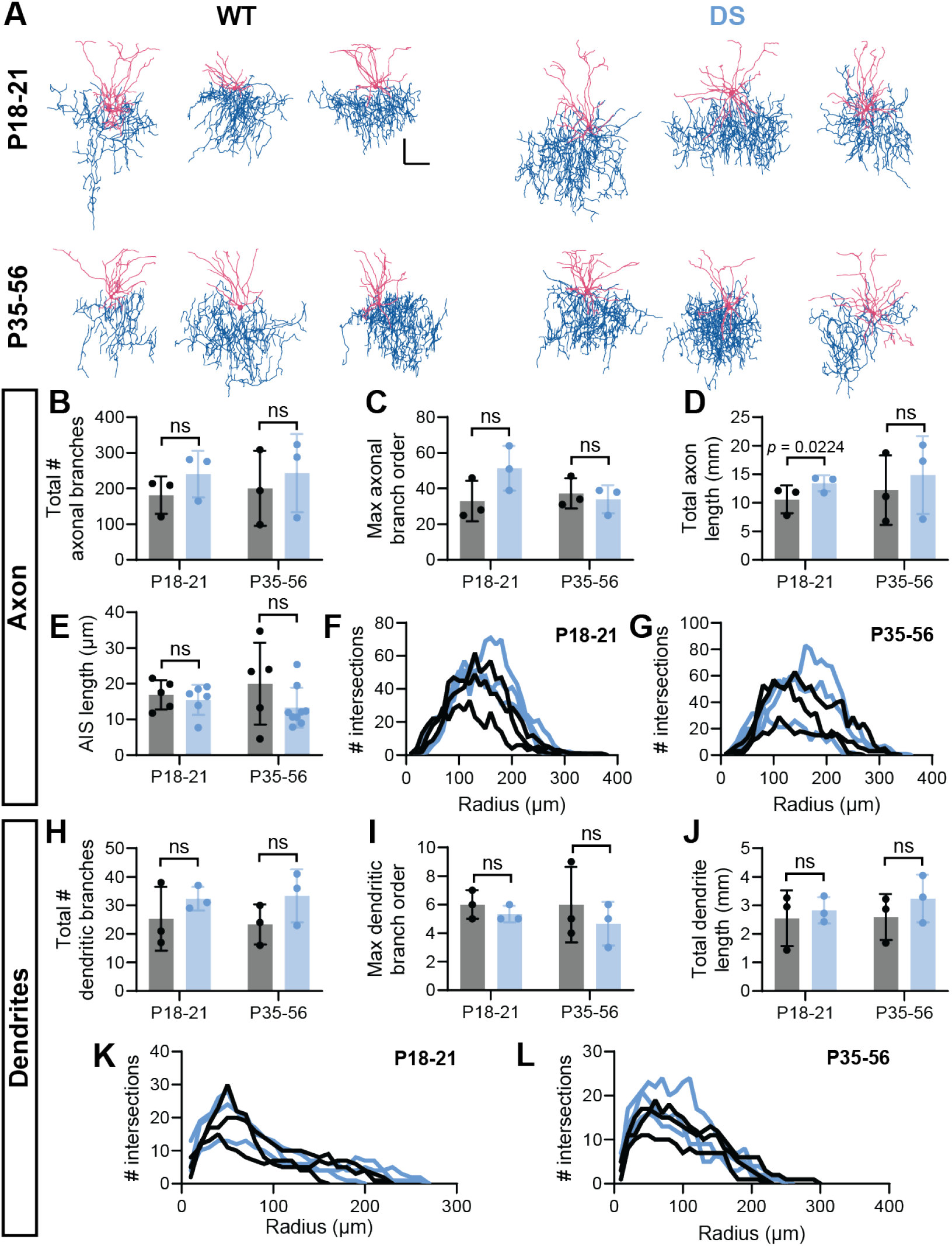
Chandelier cell morphology is not affected by loss of Scn1a. **A**, Reconstructions of WT and *Scn1a*^*+/-*^ chandelier cells (ChCs) at P18-21 or P35-56. Dendrites and soma are shown in pink; axons are shown in blue. Scale bars show 100 µm. **B-L**, Quantification of ChC axonal (B-G) and dendritic (H-L) morphology. **B and H**, Total number of axonal/dendritic branches. **C and I**, Maximal axonal/dendritic branch order. **D and J**, Total axonal/dendritic length. **E**, AIS length as measured by AnkG staining. **F and K**, Sholl analysis of axon/dendrite at P18-21 (axon, radius p = 0.0009, genotype p = 0.1713; dendrite, radius p = 0.0001, genotype p = 0.4652). **G and L**, Sholl analysis of axon/dendrite at P35-56 (axon, radius p = 0.0038, genotype p = 0.6966; dendrite, radius p < 0.0001, genotype p = 0.4424). P values indicate significance of generalized linear mixed-effects models that account for the effects of age, sex, and recording multiple cells from the same animal.

## Discussion

Here, we performed the first detailed and targeted investigation of ChC electrophysiological function and morphology in a disease model. We observed that haploinsufficiency of *Scn1a* – which causes Dravet Syndrome, the most common genetic epilepsy – reduces ChC excitability at P18-21, and that excitability largely remains impaired at P35-56. We detected no differences in the axonal or dendritic morphology of *Scn1a*^*+/-*^ ChCs at either age compared to WT. Our identification of reduced excitability in *Scn1a*^*+/-*^ ChCs also suggests that ChCs, like other inhibitory neurons, rely on Nav1.1 to generate action potentials. However, these findings are distinct from our prior observations of *Scn1a*^*+/-*^ PV+ basket cells, which exhibit impaired AP generation at P18-21 but largely restored excitability at P35-56 (14). We conclude that ChCs, unlike BCs, do not similarly compensate for Scn1a haploinsufficiency.

The study of ChCs has previously been hampered by the lack of specific labeling methods. Combinatorial approaches involving *Nkx2*.*1-CreER* or *Unc5b-CreER* and timed pre- or post-natal tamoxifen injections represent important advances, but such approaches require use of more complex mouse genetics (3, 19, 20). Other groups have used genetic meth-ods that label ChCs and BCs (such as *Nkx2*.*1-Cre*) and assessed morphology or electrophysiology to distinguish between these PV subtypes (5, 19, 21). Furlanis et al recently identified an enhancer element with high specificity for ChCs over BCs and other neurons (16). However, the specificity of the enhancer is dependent on the age and route of injection,16 as with other cell type-specific enhancers identified previously. In the present study, experimental constraints required injection in early development (P1) to facilitate labeling required to perform recording at P18-21. Via this protocol, we observed significant enrichment for ChCs, but also off-target labeling, including of pyramidal cells. This approach was suitable for patch-clamp electrophysiology experiments combined with post-hoc morphological confirmation, but perhaps might not be ideal for behavioral or largescale imaging-based approaches in vivo that require higher specificity and/or for which post-hoc confirmation of cell identify would be laborious or impossible.

The precise targeting of the pyramidal cell AIS positions ChCs as key regulators of cerebral cortex activity. However, in earlier studies, it was unclear whether ChCs played an excitatory or inhibitory role in cortical circuits. More recently, Lipkin and Bender demonstrated that iontophoretic application of GABA to the AIS reduces AP firing even when the chloride reversal potential was artificially increased to -50 mV, an effect likely due to shunting inhibition (22). Dudok et al. also demonstrated that optogenetic activation of hippocampal ChCs reduces pyramidal cell firing (20). Thus, the current evidence points to an inhibitory effect of ChCs (20–22), such that loss of ChC excitability in *Scn1a*^*+/-*^ mice likely contributes to impaired inhibition and resulting seizures/epilepsy and/or other defining features of DS.

We predicted that changes in ChC electrophysiology and in the overall excitability of the neocortex might affect morphology, but the only difference we observed was a small increase in the total axonal length in *Scn1a*^*+/-*^ ChCs at P18-21 compared to WT. Based on the lack of effect at P35-56 and the small magnitude of the relative effect, we believe this finding may be due to small sample size. ChC morphology stabilizes between P14 and P16 (21), which is around the time that the postnatal increase in *Scn1a* expression reaches a plateau (23) and that deficits in *Scn1a*^*+/-*^ BC and ChC electrophysiology are first observed (14). Thus, the morphological development of ChCs may already be complete before *Scn1a*^*+/-*^ ChCs exhibit impairment.

The deficits in *Scn1a*^*+/-*^ ChC excitability we detected at P18-21 were also present at P35-56, in contrast to our previous observations in BCs (14). There appears to be a developmentally-regulated mechanism engaged in BCs that compensates for the loss of Nav1.1. In *Scn1a*^*+/-*^ BCs, we observed an increase in the length of the AIS (14), which is thought to be a compensatory mechanism for increasing cellular excitability (24). We did not observe the same phenomenon in *Scn1a*^*+/-*^ ChCs at P35-56, which may partially explain the lack of electrophysiological normalization.

ChCs play an important role in regulating overall cerebral cortex excitability, but much remains unknown about their function. New methods of labeling ChCs will enable further study of the normal function of these cells and the potential role of ChC dysfunction in brain disorders. Our application of a new ChC-specific enhancer element uncovered a role for ChCs in the pathogenesis of Dravet Syndrome across development.

## ACKNOWLEDGEMENTS

We acknowledge the technical contributions of Dr. Xiaohong Zhang. This work was funded by a Dravet Syndrome Foundation Postdoctoral Fellowship (SFH), a Brody Family Medical Trust Fellowship from the Philadelphia Foundation (SFH), NIMH UH3MD120096 (GJF and YW), and NINDS R01 NS110869 (EMG).

## Notes

https://gin.g-node.org/GoldbergNeuroLab/Hill-et-al-2026-bioRxiv-Dysregulation-of-ChC-excitability-in-DravetSyndrome

### Competing Interest Statement

The authors have declared no competing interest.

