## Supplementary Table 1 for "Developmental dysregulation of chandelier cell excitability in a mouse model of Dravet Syndrome"

**Supplementary Table 1. Dravet Syndrome chandelier cells exhibit reduced firing frequency in adolescence.** Data are shown as mean ± standard deviation, with the number of cells indicated in parentheses. WT experiments, 13 cells were recorded from 8 mice (3F / 5M). For DS experiments, 14 cells were recorded from 6 mice (4F / 2M). *P*-values indicate the significance of generalized linear mixed-effects models that account for the effects of age, sex, and recording multiple cells from the same animal. * indicates *p* < 0.05; ** indicates *p* < 0.01. AP, action potential; AHP, afterhyperpolarization.

| Property | WT | DS | *p*-value |
| --- | --- | --- | --- |
| Resting membrane potential, mV | -64 ± 6 | -63 ± 3 | 0.3810 |
| Input resistance, MΩ | 142 ± 49 | 107 ± 24 | * 0.0488 |
| Membrane time constant, ms | 7.3 ± 1.6 | 7.4 ± 0.9 | 0.8504 |
| Membrane sag, % | 99 ± 0.4 | 99 ± 0.4 | 0.8784 |
| Maximum steady-state firing frequency, Hz | 175 ± 62 | 114 ± 20 | ** 0.0095 |
| Maximum instantaneous firing frequency, Hz | 288 ± 70 | 239 ± 36 | 0.0712 |
| Spike frequency adaptation (1^st^ vs n^th^), ratio | 0.60 ± 0.13 | 0.57 ± 0.11 | 0.4716 |
| Rheobase, pA | 163 ± 63 | 220 ± 75 | 0.0557 |
| AP threshold, mV | -30.3 ± 3.7 | -27.1 ± 4.0 | 0.0756 |
| AP half-width, ms | 0.37 ± 0.10 | 0.47 ± 0.06 | * 0.0244 |
| AP peak, mV | 28 ± 7 | 31 ± 9 | 0.3072 |
| AP amplitude, mV | 57 ± 7 | 58 ± 6 | 0.8511 |
| AHP amplitude, mV | 17 ± 3 | 16 ± 2 | 0.3225 |
| Maximum upstroke velocity, mV/ms | 351 ± 65 | 282 ± 55 | ** 0.0077 |
| Maximum downstroke velocity, mV/ms | -185 ± 67 | -135 ± 32 | * 0.0427 |
