## Supplementary Table 2 for "Developmental dysregulation of chandelier cell excitability in a mouse model of Dravet Syndrome"

**Supplementary Table 2. Dravet Syndrome chandelier cells remain impaired in early adulthood .** Data are shown as mean ± standard deviation, with the number of cells indicated in parentheses. WT experiments, 16 cells were recorded from 10 mice (5F / 5M). For DS experiments, 18 cells were recorded from 8 mice (2F / 6M). *P*-values indicate the significance of generalized linear mixed-effects models that account for the effects of age, sex, and recording multiple cells from the same animal. * indicates *p* < 0.05; ** indicates *p* < 0.01. AP, action potential; AHP, afterhyperpolarization.

| Property | WT | DS | *p*-value |
| --- | --- | --- | --- |
| Resting membrane potential, mV | -66 ± 6 | -69 ± 5 | 0.1730 |
| Input resistance, MΩ | 144 ± 34 | 155 ± 36 | 0.3952 |
| Membrane time constant, ms | 6.9 ± 1.2 | 6.8 ± 1.1 | 0.9681 |
| Membrane sag, % | 99 ± 0.6 | 100 ± 0.2 | 0.1104 |
| Maximum steady-state firing frequency, Hz | 137 ± 27 | 115 ± 35 | * 0.0482 |
| Maximum instantaneous firing frequency, Hz | 285 ± 44 | 271 ± 33 | 0.3252 |
| Spike frequency adaptation (1^st^ vs n^th^), ratio | 0.52 ± 0.10 | 0.46 ± 0.10 | 0.0905 |
| Rheobase, pA | 155 ± 41 | 148 ± 47 | 0.6146 |
| AP threshold, mV | -32.2 ± 3.2 | 32.0 ± 3.6 | 0.8696 |
| AP half-width, ms | 0.41 ± 0.07 | 0.46 ± 0.05 | * 0.0195 |
| AP peak, mV | 23 ± 9 | 19 ± 5 | 0.1471 |
| AP amplitude, mV | 55 ± 9 | 51 ± 7 | 0.1690 |
| AHP amplitude, mV | 17 ± 2 | 16 ± 3 | 0.0786 |
| Maximum upstroke velocity, mV/ms | 283 ± 66 | 231 ± 43 | ** 0.0088 |
| Maximum downstroke velocity, mV/ms | -164 ± 47 | -135 ± 32 | * 0.0360 |
